# Intravital single-molecule microscopy reveals protein kinetics in live mammalian tissues

**DOI:** 10.64898/2026.02.25.708035

**Authors:** Marco Heydecker, Desu Chen, Andrius Masedunskas, Melissa Mikolaj, Kedar Narayan, Jiji Chen, Harshad Vishwasrao, Tobias Meckel, Edna Hardeman, Peter Gunning, Roberto Weigert

## Abstract

Quantifying individual protein behavior in living mammalian tissues remains challenging. Here we introduce intravital single-molecule microscopy (iSiMM), an imaging and analysis framework that combines instant structured illumination microscopy–guided tissue navigation with highly inclined and laminated optical sheet illumination. In live mice, iSiMM enables single-emitter detection and tracking of endogenously expressed fluorescently tagged proteins at 30 ms per frame. Trajectory-based modeling quantifies diffusion coefficients and model-defined bound and free fractions, while localization-density reconstructions map molecular organization. We illustrate the approach using the actomyosin cytoskeleton, resolving isoform-specific mobility and responses to physiological stimulation and genetic perturbation. Single-molecule detection in pancreas and skeletal muscle, including after in vivo transfection, further demonstrates feasibility across additional tissues and labeling strategies. iSiMM provides a framework for investigating molecular behavior in the physiological context of living mammalian tissues.

## Introduction

Understanding how cells function within organs requires measuring not only where proteins localize, but how individual molecules move and redistribute during physiological responses. Single-molecule imaging provides access to heterogeneous molecular behavior that is obscured by ensemble fluorescence measurements, including differences in mobility within spatially overlapping populations.^**1,2**^ Yet obtaining equivalent measurements in living mammalian tissues remains challenging. Tissue architecture, extracellular interactions and mechanical constraints establish environments that are difficult to reproduce in culture, while vascularization, innervation and physiological stimulation further shape cellular responses.^**3–5**^ Connecting molecular behavior to these tissue-level influences requires measurements in the living animal.

Intravital microscopy has enabled visualization of membrane trafficking, cytoskeletal remodeling and secretion in live animals at subcellular resolution.^**4–9**^ However, changes in ensemble fluorescence intensity or recovery after photobleaching do not directly resolve the trajectories and mobility distributions of individual proteins. Extending single-molecule measurements to these settings requires sufficient signal-to-noise ratio to distinguish individual emitters against tissue background, control of motion, and an imaging strategy that preserves anatomical context while accessing defined subcellular structures. The methodological challenge is therefore to connect tissue-scale navigation and single-emitter detection with quantitative analysis of molecular behavior during physiological responses and after targeted perturbations.

Here we introduce intravital single-molecule microscopy (iSiMM), an imaging and analysis framework for measuring protein mobility and spatial organization in optically accessible regions of living mouse tissues. The workflow couples instant structured illumination microscopy (iSIM)^**10**^ for tissue navigation and region selection with highly inclined and laminated optical sheet (HILO) microscopy^**1**^ which reduces out-of-focus excitation for single-emitter detection. The two imaging paths share the same objective and tissue preparation. Fluorescent knock-in reporters provide access to endogenously expressed proteins,^**9,11,12**^ while trajectory reconstruction^**13**^, temporal localization-density mapping^**14**^ and two-state modeling^**2**^ yield complementary measurements of molecular organization, diffusion coefficients and the relative fractions of low-mobility, bound and more mobile, free populations. These populations are defined by their fitted mobility rather than by a specific biochemical interaction.

We applied iSiMM to salivary gland acinar cells, a surface-accessible epithelial system that undergoes volume expansion during β-adrenergic stimulation.^**15**^ This response requires remodeling of the basolateral plasma membrane, but how membrane architecture and cytoskeletal dynamics are coordinated in vivo has remained unclear. Previous intravital studies documented secretory granule fusion and compensatory membrane retrieval at the apical surface,^**6,9**^ without detecting bulk basolateral exocytosis during stimulation.^**5,7,9**^ These observations suggest that expansion could draw on a pre-existing membrane reservoir,_ raising the question of how the actomyosin cytoskeleton regulates its organization and deployment. Nonmuscle myosin II (NMII) and tropomyosin 3.1 (Tpm3.1), an actin-associated regulator of myosin interactions,^**16,17**^ provide a molecular system in which to connect protein behavior to the physiological membrane response.

Using iSiMM, we find that NMII trajectories outline a meshwork of actin-rich basolateral membrane domains and reveal isoform-specific differences in molecular mobility. Combining iSiMM with intravital membrane-reporter FRAP, FIB-SEM and targeted perturbations allows molecular mobility to be interpreted alongside membrane recovery, ultrastructure and physiological remodeling. During stimulation, membrane unfolding accompanies increased NMIIA mobility and a reduced bound fraction, whereas Tpm3.1 exhibits the opposite mobility response. Loss of Tpm3.1 alters the spatial organization and mobility of both NMII isoforms, identifying a regulatory relationship within this membrane-associated cytoskeleton. We further demonstrate imaging feasibility in the pancreas of fluorescent reporter mice and in skeletal muscle following in vivo transfection. Together, these results establish iSiMM as a framework for connecting single-molecule behavior to subcellular organization and physiological remodeling in living mammalian tissues.

## Results

### iSiMM combines iSIM-guided intravital imaging with HILO single-molecule detection in live mice

To enable single-molecule imaging in intact living tissue, we developed an intravital imaging workflow that couples rapid tissue navigation with shallow-angle single-molecule detection. The system was built on an inverted microscope configured with two complementary optical paths: an instant structured illumination microscopy (iSIM) path for live tissue navigation and region selection, and a TIRF–HILO path for high-contrast single-molecule imaging (Fig. 1A and Fig. S1B). iSIM provides rapid optical sectioning and volumetric imaging through multifocal excitation and image reassignment, whereas highly inclined and laminated optical sheet (HILO) illumination reduces out-of-focus excitation to enable single-molecule imaging below the tissue surface^10,18^. In this configuration, single-emitter detection of endogenously expressed fluorescently tagged proteins was facilitated by iSIM-guided selection of low-background regions, imaging of structures near the tissue surface, and HILO-mediated suppression of out-of-focus fluorescence. In the selected regions, fluorescent signals were sufficiently sparse to resolve individual emitters (Fig. 1B,C).

**Figure 1.**
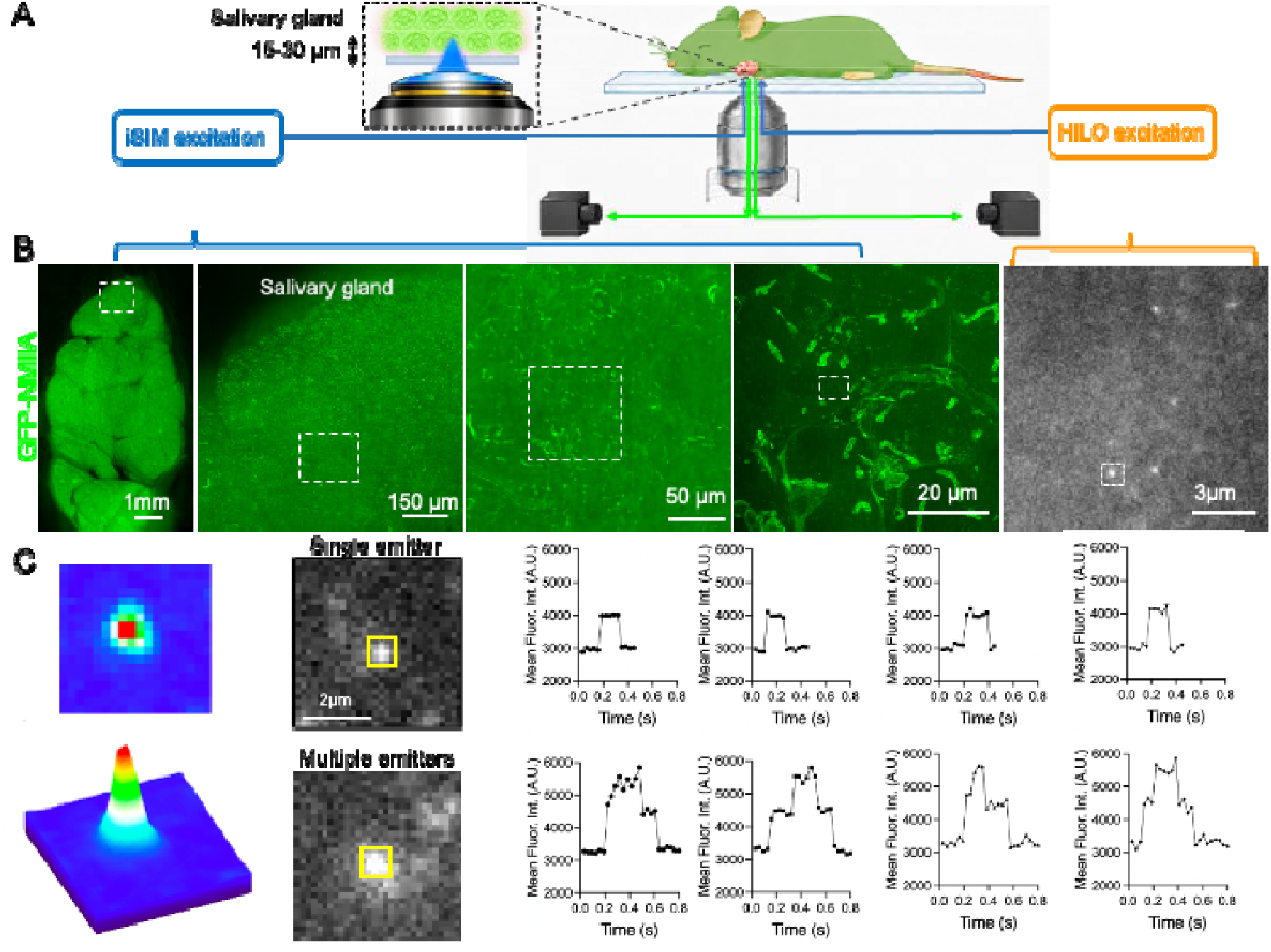
iSiMM enables intravital single-molecule imaging in live mouse salivary gland. (A) Schematic of the iSiMM workflow showing an anesthetized mouse with the surgically exposed salivary gland positioned on a coverslip above an inverted microscope objective. The system combines an iSIM path for multifocal intravital imaging and tissue navigation with a TIRF–HILO path operated in HILO mode for single-molecule imaging. A magnified side view illustrates the tissue geometry: the gland remains intact in the living animal, the coverslip contacts the exposed gland surface, acini are the first structures encountered below the surface, and the imaging plane lies approximately 15–30 μm beneath the surface. The preparation and detailed optical configuration are shown in Fig. S1A and Fig. S1B, respectively. (B) Multiscale imaging workflow in GFP–NMIIA mice. iSIM is used to image the exposed salivary gland and progressively select smaller regions of interest. The selected region is then imaged through the TIRF–HILO path, operated in HILO mode, to detect sparse single-molecule fluorescence signals. The GFP–NMIIA data shown were acquired using 488-nm excitation. See Movie S1. (C) Validation of single-emitter detection in live tissue. Representative diffraction-limited GFP–NMIIA puncta and fluorescence intensity traces distinguish single-emitter from multiple-emitter events.

The submandibular salivary gland was surgically exteriorized in anesthetized mice and stabilized against a coverslip above the inverted objective, while maintaining its vascular and neural connections (Fig. S1A). This preparation follows established intravital approaches for imaging salivary gland secretion and actomyosin remodeling.^7,10–12^ Acini are the first tissue structures encountered from the gland surface, and the basal surface of acinar cells can be imaged approximately 15–30 μm below the coverslip-contacting surface (Fig. 1A).

For the GFP–NMIIA experiments shown in Fig. 1, we used 488-nm excitation. The iSIM path was used first to navigate from the exposed gland to progressively smaller regions within the tissue and identify fields suitable for single-molecule acquisition by avoiding migratory cells and high background areas. The selected region was then imaged using the TIRF–HILO path, operated in HILO mode, to detect sparse GFP–NMIIA fluorescence signals in the living animal (Fig. 1B; Movie S1). The two imaging modalities shared the same objective and tissue preparation but used dedicated illumination and detection paths (Fig. S1B).

Using GFP–NMIIA knock-in mice,^19,20^ individual fluorescent puncta were detected in selected regions of the exposed salivary gland. To validate that these puncta corresponded to single fluorescent emitters, we analyzed their spatial and temporal fluorescence properties. Representative GFP–NMIIA puncta showed diffraction-limited intensity profiles compatible with single emitters, and fluorescence traces displayed stepwise intensity changes consistent with single- and multiple-emitter events (Fig. 1C). These data establish that iSiMM can detect endogenously expressed fluorescent molecules in a living mouse and provide the basis for subsequent tracking and kinetic analysis.

### iSiMM reveals spatially organized myosin II dynamics at the basolateral plasma membrane

Having established single-emitter detection in living tissue, we next used single-particle tracking to examine the spatial organization and mobility of endogenous nonmuscle myosin II (NMII). In GFP–NMIIA and GFP–NMIIB knock-in mice,^11,12^ images were acquired at 30 ms per frame, and individual fluorescent emitters were linked into trajectories using TrackMate^13^ (Fig. 2A). Overlays of trajectories from 10,000-frame acquisitions revealed that molecular motion was not uniformly distributed across the imaged cellular area. Instead, trajectories preferentially populated discrete regions that collectively outlined a meshwork-like pattern, with multiple trajectories occupying the same regions for both NMII isoforms (Fig. 2B). These observations suggested that NMII dynamics were organized around a defined subcellular structure.

**Figure 2.**
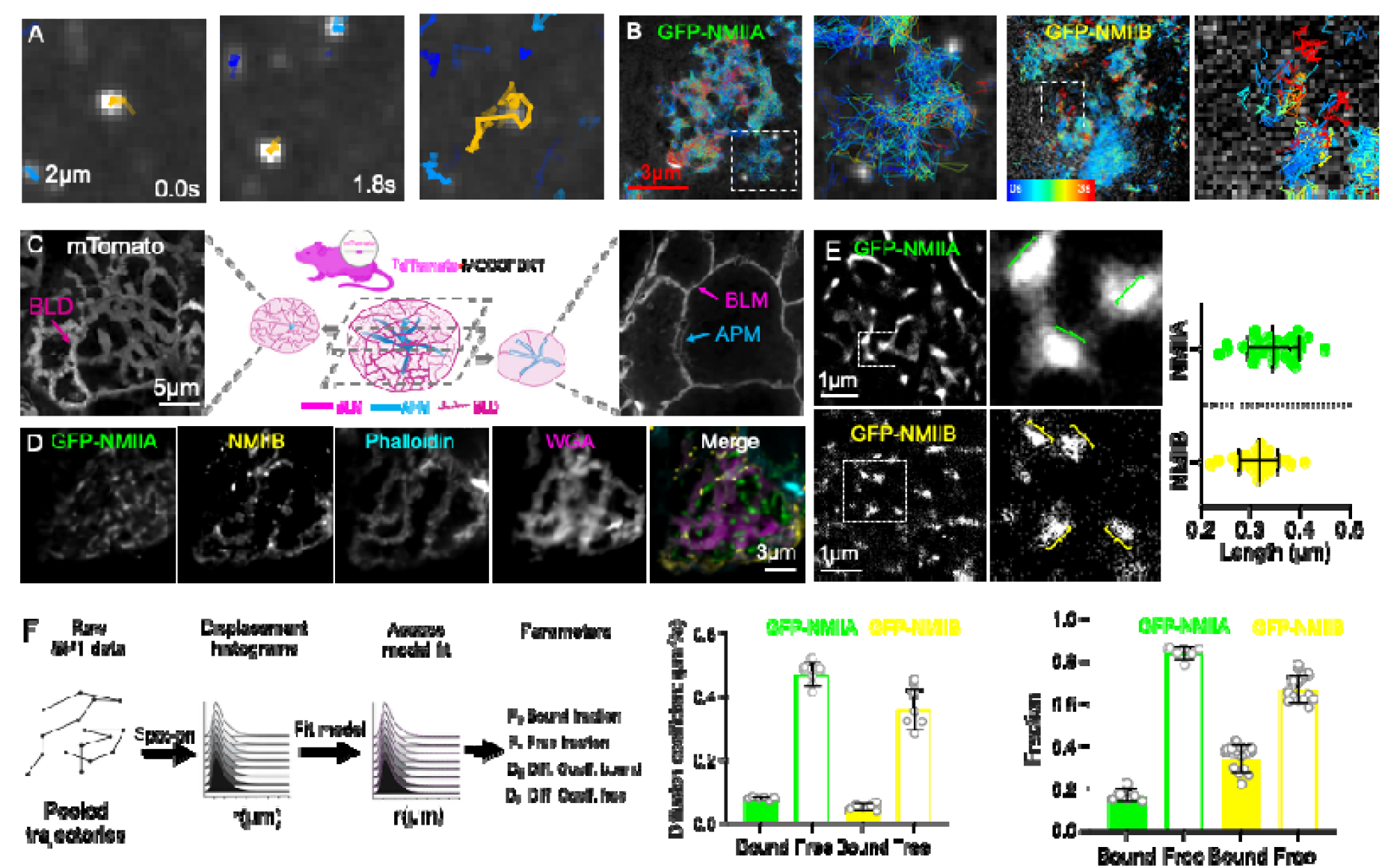
Single-molecule tracking reveals the spatial organization and mobility of nonmuscle myosin II at basolateral membrane domains. **(A)** Representative tracking of an individual GFP–NMIIA molecule by intravital single-molecule microscopy (iSiMM) in a live mouse salivary gland. Images show the tracked molecule at the initial and final time points, with its reconstructed trajectory highlighted in yellow. Scale bar, 2 μm. **(B)** Overlays of GFP–NMIIA and GFP–NMIIB trajectories reconstructed from 10,000-frame iSiMM acquisitions. Trajectories are color-coded by track duration, as indicated by the color key. Enlargements of the boxed regions show multiple trajectories occupying spatially restricted areas within a meshwork-like pattern. Scale bar, 3 μm. **(C)** Identification of basolateral membrane domains using the membrane-targeted tdTomato reporter, mTom. Left, representative intravital optical section showing the membrane meshwork at the basal surface of an acinar cell. Center, schematic illustrating the membrane reporter and the relationship between the imaged domains and acinar membrane architecture. Right, an optical section showing the basolateral plasma membrane (BLM) and apical plasma membrane (APM). BLD, basolateral membrane domain. Scale bar, 5 μm, left. **(D)** Fluorescence imaging of fixed salivary gland tissue from GFP–NMIIA mice showing GFP–NMIIA (green), NMIIB (yellow), F-actin labeled with phalloidin (cyan), and WGA (magenta). The merged image shows the association of both NMII isoforms with the actin-rich, WGA-positive membrane meshwork. Individual channels are displayed in grayscale. Scale bar, 3 μm. **(E)** ThunderSTORM localization-density reconstructions generated from 10,000-frame iSiMM datasets of GFP–NMIIA and GFP–NMIIB. Enlargements of the boxed regions show elongated localization densities. Colored line segments indicate representative length measurements for NMIIA (green) and NMIIB (yellow). The graph shows the lengths of individual localization densities, with each point representing one measured density; *n* = 31 densities for each isoform. Lengths were determined from line profiles extending to the return of the signal to baseline. Data are mean ± SD. Scale bars, 1 μm, overview images. **(F)** Left, schematic of the Spot-On analysis workflow. Displacement distributions calculated from pooled single-particle trajectories are fitted using a two-state model to estimate the bound and free fractions, *F*_B_ and *F*_F_, and their corresponding diffusion coefficients, *D*_B_ and *D*_F_. Right, diffusion coefficients and population fractions for GFP–NMIIA and GFP–NMIIB. Bound and free populations are defined by their fitted mobility. GFP–NMIIA bound and free fractions are 0.16 ± 0.03 and 0.84 ± 0.03, respectively. GFP–NMIIA: *N* = 3 mice, 6 cells, 391,711 trajectories; GFP–NMIIB: *N* = 3 mice, 6 cells, 361,614 trajectories. Data are mean ± SD. Spot-On fitting was performed separately for each cell. Cell-level diffusion-coefficient estimates were averaged within each animal, and NMIIA and NMIIB were compared using unpaired two-sided Welch’s t-tests on animal means. Individual-cell values are shown to illustrate within-animal variability and were not used as independent replicates for statistical inference. Statistical significance is indicated for the comparisons shown: ns, *P* > 0.05; **, *P* ≤ 0.01; ***, *P* ≤ 0.001; ****, *P* ≤ 0.0001

To identify the structure underlying this pattern, we examined acinar membrane organization in mice expressing membrane-targeted tdTomato (mTom).^19^ Intravital imaging revealed a similarly organized meshwork of membrane-enriched regions at the basal surface of acinar cells. Imaging of the broader acinar architecture placed these regions within the basolateral plasma membrane, distinct from the apical membrane network (Fig. 2C). We refer to these regions as basolateral membrane domains (BLDs). Complementary labeling supported their membrane identity: wheat germ agglutinin (WGA), a plasma membrane marker,^20^ overlapped with mTom-positive domains, whereas in wild-type glands lacking the membrane reporter, WGA labeling overlapped with the basolateral marker Na^+^/K^+^-ATPase^21^ (Fig. S2A).

We then examined the relationship between NMII, F-actin and the membrane meshwork. Imaging of fixed glands from GFP–NMIIA mice showed that both GFP–NMIIA and immunolabeled NMIIB were enriched along WGA-positive, actin-rich BLDs (Fig. 2D). Supporting analyses showed colocalization of F-actin with mTom-positive domains and positive Pearson’s correlations between F-actin and either NMII isoform (Fig. S2B). The lower correlations between NMII and F-actin, compared with those between mTom and F-actin, were consistent with association within shared domains but nonidentical spatial distributions. Together, these findings associated the meshwork revealed by single-molecule tracking with actin-rich regions of the basolateral plasma membrane.

To resolve NMII organization within these domains, we generated super-resolution localization-density reconstructions using ThunderSTORM.^14^ Reconstructions from 10,000-frame datasets revealed discrete, high-density sites of NMII localization distributed within the membrane-associated meshwork (Fig. 2E). Many of these densities were elongated, with measured lengths on the order of 0.3 μm for both isoforms. These dimensions were compatible with reported NMII minifilament lengths,^22^ suggesting that the time-integrated localization patterns captured sites of higher-order NMII organization in vivo.

We next quantified molecular mobility using the two-state Spot-On framework, which fits displacement distributions to estimate diffusion coefficients and the relative fractions of low-mobility, bound and more mobile, free populations.^2^ GFP–NMIIA was predominantly assigned to the free population, with bound and free fractions of 0.16 ± 0.03 and 0.84 ± 0.03, respectively (mean ± SD; Fig. 2F). NMIIB was also predominantly free but exhibited a larger bound fraction and lower diffusion coefficients in both fitted states than NMIIA. Here, bound and free denote mobility states inferred by the model. Thus, although NMIIA and NMIIB were associated with the same membrane meshwork, they exhibited different mobility and state-occupancy profiles. These results connect single-molecule trajectories to defined membrane-associated structures and establish a quantitative baseline for examining how endogenous NMII dynamics respond to physiological stimulation.

### NMII stabilizes folded basolateral membrane reservoirs that remodel during secretion

The enrichment of F-actin and NMII within BLDs suggested that the actomyosin cytoskeleton could regulate basolateral membrane remodeling during secretion. Salivary gland acinar cells expand during β-adrenergic stimulation,^15^ providing an opportunity to relate changes in membrane architecture to NMII behavior in vivo. Following stimulation with isoproterenol (ISOP), intravital volumetric analysis of mTom-expressing glands revealed an approximately 15% increase in acinar volume over 30 min, accompanied by widening of BLDs (Fig. 3A). Representative images with overlaid boundaries further illustrated the expansion after stimulation (Fig. S3A).

**Figure 3.**
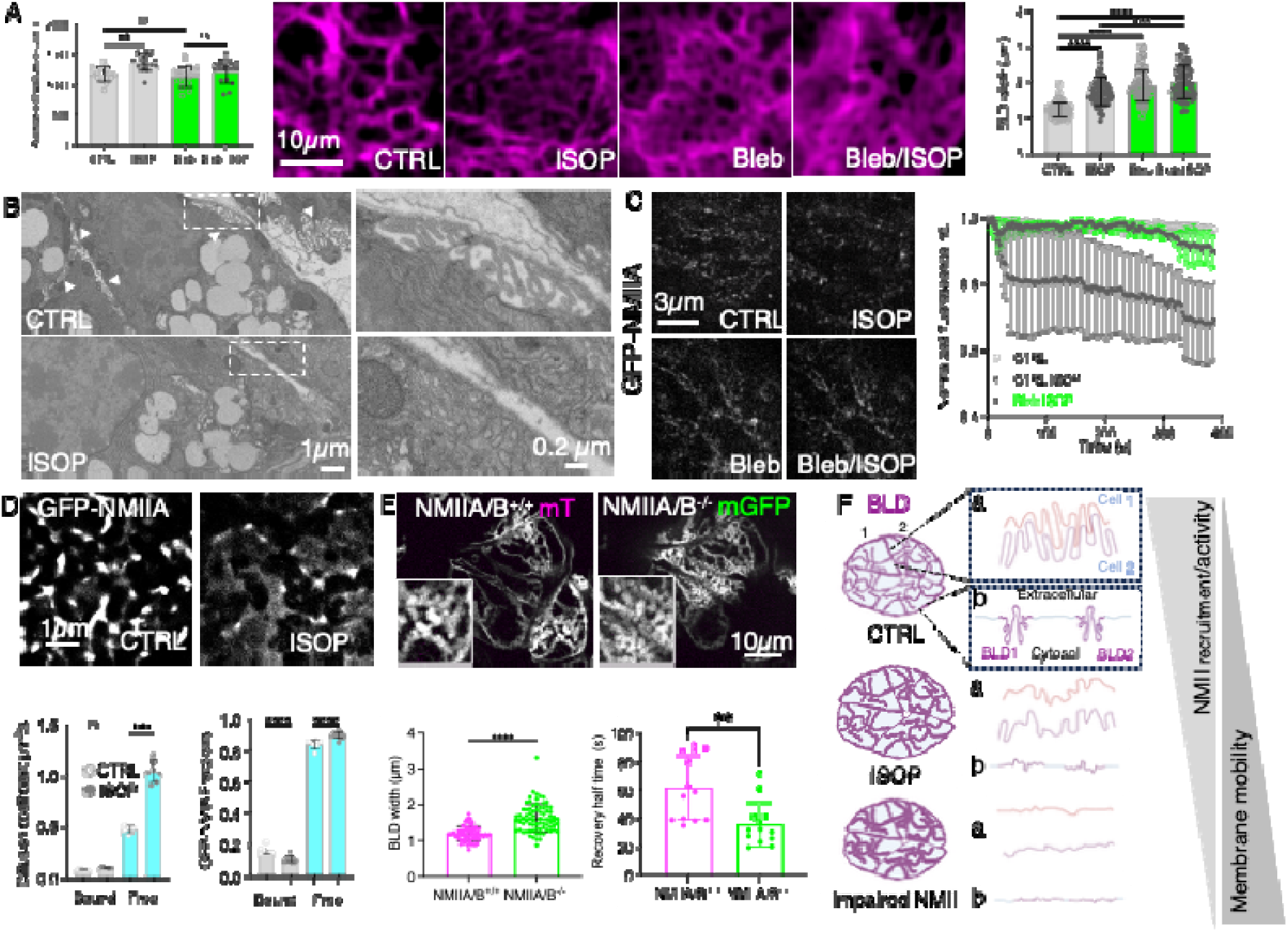
NMII-dependent remodeling of basolateral membrane folds during acinar cell expansion. (A) Effects of β-adrenergic stimulation and NMII inhibition on acinar cell volume and basolateral membrane domain (BLD) width in mTom-expressing mice. Left, quantification of acinar cell volume under control conditions (CTRL), after isoproterenol stimulation (ISOP), after treatment with para-nitroblebbistatin (Bleb), and after combined inhibitor treatment and stimulation (Bleb/ISOP). Center, representative intravital optical sections showing mTom-labeled basolateral membranes under the four conditions. Right, quantification of BLD width. Volume analysis: *N* = 3 mice, 15 acini per condition. Two-way ANOVA test was used to test for significance. Scale bar, 10 μm. (B) Representative FIB-SEM images of salivary gland acini under control and ISOP-stimulated conditions. Arrowheads indicate regions of folded basolateral membrane. Boxed regions are enlarged on the right, showing closely apposed membrane folds under control conditions and reduced folding after stimulation. Scale bars, 1 μm, overview images; 0.2 μm, enlargements. (C) Intravital subcellular imaging of GFP–NMIIA at the basolateral membrane under CTRL, ISOP, Bleb and Bleb/ISOP conditions. The graph shows normalized GFP–NMIIA fluorescence over time in unstimulated controls, ISOP-stimulated controls and ISOP-stimulated glands pretreated with para-nitroblebbistatin. Inhibitor treatment attenuates the stimulation-associated decrease in GFP–NMIIA fluorescence. *N* = 3 mice, 6 cells per condition. Scale bar, 3 μm. (D) ThunderSTORM localization-density reconstructions of GFP–NMIIA generated from 10,000-frame iSiMM acquisitions under control and ISOP-stimulated conditions. Graphs show diffusion coefficients and population fractions obtained using a two-state Spot-On model. ISOP increases the diffusion coefficient of the free population and decreases the bound fraction, with a corresponding increase in the free fraction. The diffusion coefficient of the bound population does not change significantly. Bound and free denote mobility states inferred by the model. *N* = 3 mice, 6 cells per condition. Cell-level diffusion-coefficient estimates were averaged within each animal, and CTRL and ISOP were compared using unpaired two-sided Welch’s t-tests on animal means. Individual-cell values are shown to illustrate within-animal variability and were not used as independent replicates for statistical inference. Scale bar, 1 μm. (E) Basolateral membrane organization and membrane-reporter recovery in conditional NMIIA/B-deficient acinar cells and neighboring controls. Control cells are identified by membrane-targeted tdTomato expression (mT), whereas recombined NMIIA/B-deficient cells express membrane-targeted GFP (mGFP). Membrane channels are displayed in grayscale; insets show enlarged views of BLDs. Graphs show BLD width and fluorescence recovery half-times measured by FRAP. NMIIA/B-deficient cells exhibit wider BLDs and faster membrane-reporter recovery. *N* = 3 mice, 12 cells per condition. Statistical analyses are described in the Methods. Scale bar, 10 μm. (F) Model of NMII-dependent regulation of basolateral membrane folding. Schematics illustrate membrane organization at cell–cell interfaces (a) and at the basal cell surface (b) under resting conditions, after ISOP stimulation and following impairment of NMII function. Reduced NMII recruitment or activity is proposed to relax constraints on folded membrane architecture and increase membrane mobility. Data are presented as mean ± SD. Statistical significance is indicated for the comparisons shown: ns, *P* > 0.05; **, *P* ≤ 0.01; ***, *P* ≤ 0.001; ****, *P* ≤ 0.0001.

Inhibition of NMII with para-nitroblebbistatin^23^ increased BLD width even in the absence of stimulation but markedly attenuated ISOP-induced acinar expansion (Fig. 3A and Fig. S3A). Thus, broadening of the membrane domains was not sufficient to reproduce the coordinated volume increase elicited by stimulation. These findings suggested that NMII contributes both to maintaining basolateral membrane organization under resting conditions and to its regulated remodeling during secretion.

To determine the structural basis of this remodeling, we examined basolateral membrane ultrastructure using focused ion beam–scanning electron microscopy (FIB-SEM).^24^ In unstimulated glands, the basolateral membrane formed deep, closely apposed folds at cell–cell interfaces and along the peripheral surface of acini. Following ISOP stimulation, these folds were markedly reduced, revealing a less convoluted membrane organization (Fig. 3B). These observations support the interpretation that BLDs contain a pre-existing membrane reservoir that unfolds during acinar expansion, providing additional surface area without requiring equivalent membrane addition.^25^

We next asked whether membrane unfolding was accompanied by changes in NMII organization and molecular mobility. Ensemble intravital imaging showed a rapid decrease in GFP–NMIIA fluorescence at BLDs after ISOP stimulation, whereas fluorescence remained relatively stable in unstimulated controls. The stimulation-associated decrease was attenuated by para-nitroblebbistatin treatment (Fig. 3C), linking changes in local NMIIA enrichment to the NMII-dependent remodeling response.

To resolve the molecular behavior accompanying this redistribution, we applied iSiMM to GFP–NMIIA before and after stimulation. ThunderSTORM reconstructions^14^ generated from 10,000-frame acquisitions showed that the discrete, high-density localization sites observed under basal conditions became more dispersed following ISOP treatment (Fig. 3D). Two-state Spot-On analysis^2^ revealed a marked increase in the diffusion coefficient of the free population, whereas the diffusion coefficient of the bound population did not change significantly. Stimulation also produced a modest decrease in the bound fraction and a corresponding increase in the free fraction (Fig. 3D). Here, bound and free denote mobility states inferred by the model. Thus, membrane unfolding was accompanied by reduced local NMIIA enrichment and a shift toward a more mobile molecular population.

To test whether NMII is required to maintain basolateral membrane organization, we examined acinar cells conditionally deficient in both NMIIA and NMIIB. Mist1CreERT2-mediated recombination of floxed NMIIA/B alleles^26–28^ was coupled to the mTom/mGFP membrane reporter,^19^ allowing recombined, mGFP-positive NMIIA/B-deficient cells to be distinguished from neighboring mTom-positive controls within mosaic glands (Fig. S3C). Loss of NMIIA/B increased BLD width, recapitulating the broadening observed after pharmacological inhibition (Fig. 3E).

We then used fluorescence recovery after photobleaching (FRAP) to assess membrane-reporter recovery in NMIIA/B-deficient and neighboring control cells. Representative image sequences showed recovery of the membrane-targeted fluorescent reporters within the bleached regions (Fig. S3C), and quantitative analysis revealed shorter recovery half-times in NMIIA/B-deficient cells than in neighboring controls (Fig. 3E). In addition to the increased BLD width, this result is consistent with reduced stabilization of membrane-associated domains in the absence of NMII.

These results support a model in which NMII stabilizes folded basolateral membrane reservoirs under resting conditions. During secretion, reduced NMIIA enrichment and increased molecular mobility accompany relaxation and unfolding of these reservoirs as acini expand. Inhibition or loss of NMII disrupts membrane organization, but inhibition does not reproduce the coordinated volume expansion elicited by stimulation (Fig. 3F). Together, single-molecule tracking, intravital membrane-reporter FRAP and FIB-SEM provide complementary molecular, ensemble and ultrastructural readouts that connect NMII behavior to physiological membrane remodeling.

### Tpm3.1 regulates NMII molecular mobility and organization at basolateral membrane domains

The coupling of NMII dynamics to basolateral membrane remodeling prompted us to investigate how these dynamics are regulated within BLDs. We focused on tropomyosin 3.1 (Tpm3.1), an actin-associated protein implicated in the organization of functionally distinct actin filaments and their interactions with myosin II.^16,17^ Fluorescence imaging of endogenous mNeonGreen-tagged Tpm3.1^9^ together with LifeAct–RFP-labeled F-actin showed that Tpm3.1 was enriched along the actin-rich meshwork of BLDs (Fig. 4A), positioning it as a candidate regulator of the NMII behavior associated with membrane remodeling. We first asked whether Tpm3.1 dynamics changed during the same physiological response that increased NMIIA mobility. Using iSiMM, we tracked individual Tpm3.1–mNeonGreen molecules in live salivary glands before and after ISOP stimulation. ThunderSTORM localization-density reconstructions^14^ generated from 10,000-frame acquisitions showed that Tpm3.1 remained associated with the BLD meshwork after stimulation (Fig. 4B). However, two-state Spot-On analysis^2^ revealed a marked reduction in the diffusion coefficient of the free population, together with an increased bound fraction and a corresponding decrease in the free fraction. The diffusion coefficient of the bound population did not change significantly. Thus, whereas stimulation increased NMIIA mobility and reduced its bound fraction (Fig. 3D), it shifted Tpm3.1 toward greater occupancy of the low-mobility, bound state (Fig. 4B). These contrasting responses indicate that physiological stimulation differentially regulates the molecular behavior of components within the same membrane-associated cytoskeletal network.

**Figure 4.**
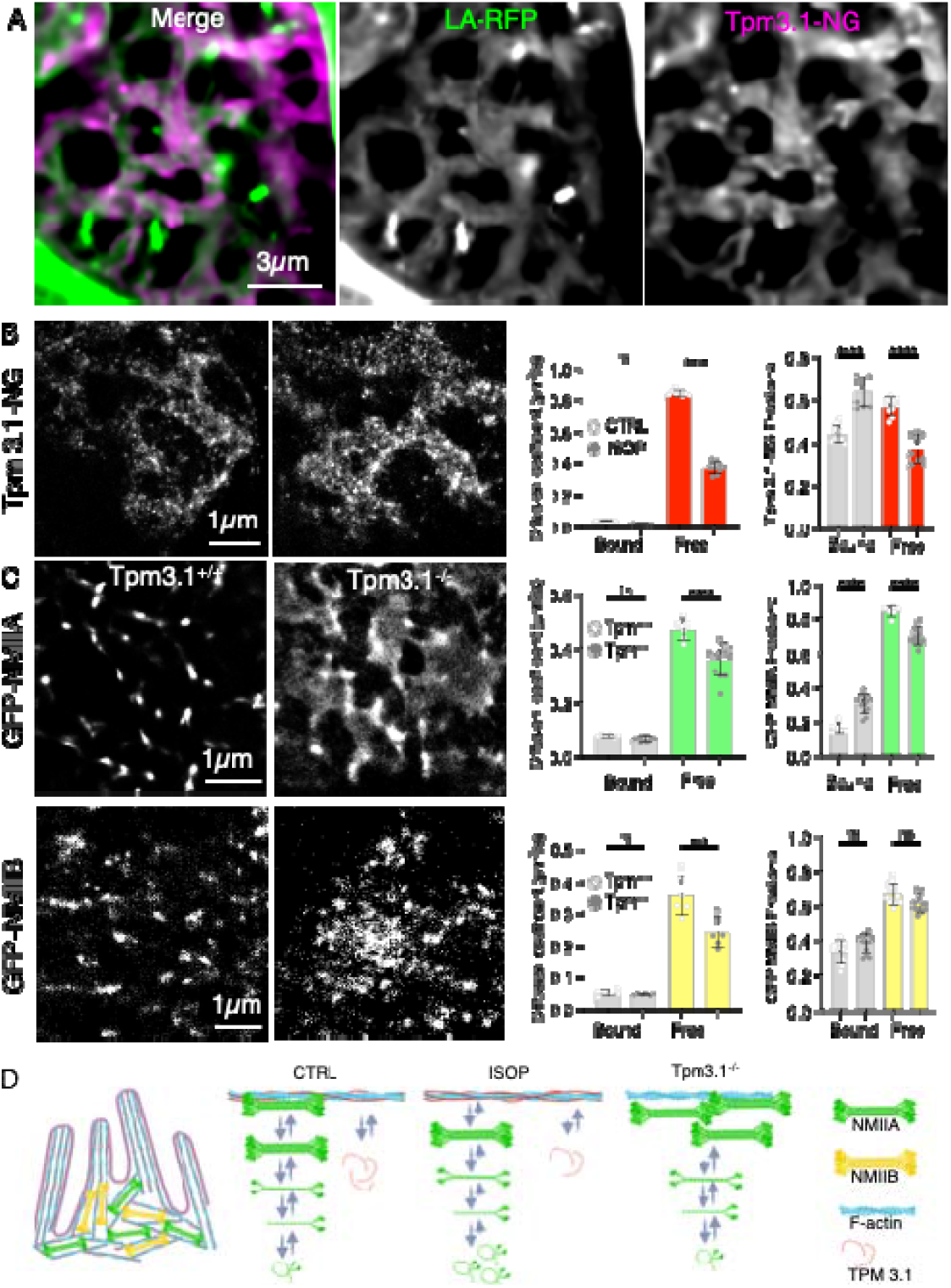
Tpm3.1 regulates NMII molecular mobility and spatial organization at basolateral membrane domains. **(A)** Fluorescence images showing the spatial relationship between Tpm3.1–mNeonGreen (Tpm3.1–NG) and LifeAct–RFP-labeled F-actin (LA-RFP) in salivary gland acinar cells. The merged image displays Tpm3.1–NG in magenta and LifeAct–RFP in green, showing their enrichment within the basolateral membrane domain (BLD) meshwork. Individual channels are displayed in grayscale. Scale bar, 3 μm. **(B)** Left, ThunderSTORM localization-density reconstructions generated from 10,000-frame iSiMM acquisitions of Tpm3.1–NG under control conditions (left image) and after isoproterenol stimulation (ISOP; right image). Right, diffusion coefficients and population fractions estimated using a two-state Spot-On model. ISOP reduces the diffusion coefficient of the free population and increases the bound fraction, with a corresponding decrease in the free fraction. The diffusion coefficient of the bound population does not change significantly. *N* = 3 mice, 9 cells per condition; 731,941 trajectories in total. Scale bar, 1 μm. **(C)** ThunderSTORM localization-density reconstructions of GFP–NMIIA (top) and GFP–NMIIB (bottom) in control (Tpm3.1^+/+^) and Tpm3.1-deficient (Tpm3.1^−/−^) acinar cells, generated from 10,000-frame iSiMM acquisitions. Graphs show the corresponding diffusion coefficients and bound/free population fractions obtained by Spot-On analysis. Loss of Tpm3.1 reduces the diffusion coefficient of the free population for both NMII isoforms, without significant changes in their bound-state diffusion coefficients. The NMIIA bound fraction increases and its free fraction decreases, whereas changes in the NMIIB population fractions are not significant. GFP–NMIIA: *N* = 5 mice, 9 cells per condition, 1,235,594 trajectories in total. GFP–NMIIB: *N* = 5 mice, 9 cells per condition, 925,136 trajectories in total. Scale bars, 1 μm. **(D)** Model of Tpm3.1-dependent regulation of actomyosin organization at basolateral membrane folds. Left, schematic illustrating the proposed arrangement of actin and NMII around folded membranes. Right, proposed relationships between NMII assembly and molecular exchange under control, ISOP-stimulated and Tpm3.1-deficient conditions. Stimulation is associated with increased Tpm3.1 bound-state occupancy and increased NMIIA mobility, whereas Tpm3.1 deficiency reduces NMII mobility and increases NMIIA bound-state occupancy. NMIIA, green; NMIIB, yellow; F-actin, cyan; Tpm3.1, red. Bound and free denote mobility states inferred by the two-state model. Cell-level diffusion-coefficient estimates were averaged within each animal, and CTRL and ISOP were compared using unpaired two-sided Welch’s t-tests on animal means. Individual-cell values are shown to illustrate within-animal variability and were not used as independent replicates for statistical inference. Data are mean ± SD. Statistical significance is indicated for the comparisons shown: ns, *P* > 0.05; ***, *P* ≤ 0.001; ****, *P* ≤ 0.0001. Statistical analyses are described in the Methods

To determine whether Tpm3.1 contributes to NMII organization and dynamics, we next examined GFP–NMIIA and GFP–NMIIB in Tpm3.1-deficient acinar cells. Under basal conditions, loss of Tpm3.1 altered the spatial enrichment of both isoforms. The discrete, elongated NMIIA localization densities observed in controls became less clearly resolved, while NMIIB displayed a more irregular pattern of local enrichment (Fig. 4C). Spot-On analysis revealed a reduction in the diffusion coefficient of the free population for both isoforms, without significant changes in their bound-state diffusion coefficients. Tpm3.1 deficiency also increased the NMIIA bound fraction and decreased its free fraction. By contrast, the corresponding changes in NMIIB population fractions were not statistically significant (Fig. 4C). These findings establish that Tpm3.1 contributes to the spatial organization and molecular mobility of both NMII isoforms, with a detectable shift in bound-state occupancy for NMIIA.

Together, these results support a model in which Tpm3.1 regulates the balance of NMII mobility and state occupancy at membrane-associated actin structures, rather than simply promoting stable NMII recruitment (Fig. 4D). During stimulation, increased Tpm3.1 bound-state occupancy coincides with increased NMIIA mobility; conversely, Tpm3.1 loss reduces the mobility of both NMII isoforms and shifts NMIIA toward the bound population. In the context of the NMII-dependent membrane remodeling described above, these findings identify Tpm3.1 as a regulator of the actomyosin dynamics associated with the folded basolateral membrane reservoir. They also extend iSiMM from measuring physiological changes in individual proteins to resolving regulatory relationships between endogenous cytoskeletal components in living tissue.

Finally, we assessed the feasibility of single-molecule detection beyond the salivary gland, using pancreatic tissue from fluorescent reporter mice and skeletal muscle expressing Tpm3.1–mNeonGreen following in vivo transfection (Fig. S4A,B). In both preparations, imaging was performed at the tissue surface and 3 μm below it, demonstrating the feasibility of applying the approach in additional tissue contexts. These experiments also establish compatibility with alternative labeling strategies, allowing fluorescently tagged proteins to be investigated without requiring a dedicated reporter mouse line. Together with the quantitative single-molecule analyses in salivary glands, these findings establish iSiMM as an adaptable framework for connecting molecular mobility and spatial organization to physiological processes in living tissues.

## Discussion

A major gap between single-molecule biophysics and intravital cell biology has been the inability to measure molecular behavior directly within intact mammalian tissues. Single-molecule approaches have transformed our understanding of protein mobility, binding and state transitions in cultured cells and reconstituted systems, whereas intravital microscopy has revealed cellular and subcellular dynamics in their native physiological context. Until now, however, these two scales have remained largely separate. Here, iSiMM bridges that gap by enabling detection and tracking of individual fluorescently tagged proteins in living mouse tissues and by linking those measurements to defined subcellular structures and physiological responses. The combination of iSIM-guided navigation, HILO illumination, single-particle tracking, localization-density reconstruction and kinetic modeling provides a practical framework for moving from tissue-scale context to molecular-scale measurements within the same preparation.

The significance of iSiMM is therefore not simply that it detects single fluorescent emitters in vivo, but that it allows molecular behavior to be quantified in the native tissue environment. This distinction is important because ensemble intravital measurements can reveal changes in localization or fluorescence intensity, but cannot distinguish whether those changes arise from altered diffusion, redistribution among molecular states, or changes in local occupancy. In contrast, iSiMM provides measurements of molecular mobility and state fractions while preserving information about where those molecules reside within the tissue. In the present study, this was sufficient to distinguish the behavior of NMIIA and NMIIB even though both isoforms were associated with the same basolateral membrane meshwork. Detection of a single GFP emitter does not establish whether the labeled NMII heavy chain belongs to an individual holoenzyme or a higher-order assembly containing additional non-emitting fluorophores. Multistep intensity traces are consistent with multiple emitters within a diffraction-limited region but do not, by themselves, demonstrate filament assembly over time. The method therefore offers access to molecular heterogeneity that would remain hidden in conventional intravital fluorescence imaging.

An important strength of iSiMM is its complementarity with measurements that connect molecular behavior to cellular organization. Here, single-molecule measurements of protein mobility and state occupancy were interpreted alongside intravital FRAP of membrane reporters and FIB-SEM of fixed tissue. These approaches provide distinct readouts of molecular motion, ensemble fluorescence recovery and membrane ultrastructure. Together with physiological stimulation and targeted perturbations, they place molecular behavior within a structural and functional context. This combination illustrates how iSiMM can extend in vivo biophysics to the single-molecule scale and support investigation of how molecular regulation contributes to tissue function. Applied to the actomyosin cytoskeleton, this multiscale approach connects NMII molecular behavior to physiological membrane remodeling. During β-adrenergic stimulation, acinar expansion was accompanied by unfolding of basolateral membrane folds and a change in NMIIA molecular behavior, including increased mobility of the free population and reduced bound-state occupancy. Pharmacological and genetic perturbations further supported a role for NMII in maintaining the resting organization of these membrane domains. Together, these observations suggest that NMII participates in balancing membrane stabilization with the ability to remodel rapidly during secretion. Rather than acting simply as a static cortical scaffold, NMII appears to occupy a dynamically regulated molecular state whose properties change as the membrane reservoir is deployed. This provides a direct molecular link between actomyosin behavior and membrane plasticity in vivo.

The Tpm3.1 experiments extend this concept by showing that iSiMM can also be used to interrogate regulatory relationships between endogenous cytoskeletal components. Tpm3.1 and NMIIA responded differently to stimulation, and loss of Tpm3.1 altered both the spatial organization and mobility of NMII isoforms. These findings argue that remodeling of the membrane-associated cytoskeleton is not driven by uniform mobilization of all components, but by differential regulation of molecular states within a shared network. More importantly from a methodological standpoint, this demonstrates that iSiMM can move beyond descriptive measurements of a single protein to examine how one molecular component influences the behavior of another inside intact tissue.

There are important limitations to the current implementation. iSiMM is presently best suited to optically accessible regions near the surface of exposed tissues, and its performance depends on tissue stability, signal-to-background ratio and reporter behavior. In addition, the bound and free populations obtained from two-state modeling represent mobility states rather than direct measurements of biochemical binding or residence time. These parameters therefore need to be interpreted alongside complementary structural and functional measurements. Nonetheless, these limitations define the current experimental window rather than the conceptual scope of the method.

Single-molecule detection in pancreas and skeletal muscle, including after in vivo transfection, supports the feasibility of applying the imaging strategy beyond the salivary gland and across labeling approaches. These observations provide a foundation for further quantitative validation in additional tissues. With appropriate tissue access and labeling, the same framework could in principle be applied to a wide range of biological questions in which molecular behavior is shaped by the native tissue environment. Potential applications include receptor signaling, cytoskeletal remodeling, cell–cell and cell–matrix interactions, membrane trafficking, immune-cell contacts, tumor–stroma interfaces and vascular biology. In each of these settings, the central advantage would be the same: the ability to measure how individual proteins behave while the cell remains embedded in its physiological context.

More broadly, iSiMM provides a route toward integrating molecular biophysics with systems-level cell biology in vivo. Many cellular processes are ultimately governed by changes in molecular mobility, assembly and state occupancy, yet these parameters are usually measured outside the tissue environment in which they acquire physiological meaning. By bringing single-molecule measurements into living organs, iSiMM creates the possibility of asking not only where proteins are and how much of them is present, but how their molecular behavior changes during normal physiology, disease progression or therapeutic intervention. The NMII/Tpm3.1 system described here provides one example of this principle, linking molecular-state changes to membrane remodeling during secretion. The broader opportunity is to use the same approach to uncover molecular mechanisms across diverse biological systems directly in vivo.

## Materials and Methods

### Animal strains and procedures

All animal experiments were conducted in accordance with guidelines from the National Cancer Institute (NIH, Bethesda, MD, USA) and the NSW Animal Research Act (1985) and approved by the respective institutional animal care and ethics committees. Male and female mice (8–24 weeks old) were maintained on standard chow and water ad libitum.

The mT/mG Cre reporter mouse was obtained from Jackson Laboratory. Hemizygous LifeAct-GFP and LifeAct-RFP mice were provided by R. Wedlich-Soldner ^29^. GFP–NMIIA knock-in mice were generated as described ^11^ and crossed with LifeAct-RFP mice. The endogenous Tpm3.1–mNeonGreen knock-in mouse line [B6-Tpm3tm5(mNeonGreen)Hrd] was generated as described ^9^. Conditional NMIIA/B floxed mice (NMIIA/Bfl/fl) ^26,27^ were crossed with Mist1CreERT2 mice ^28^ and the mT/mG reporter strain. Cre recombination was induced by two intraperitoneal injections of tamoxifen (75 mg/kg body weight) administered two days apart. Mice were imaged 3–8 weeks after induction. Recombined (GFP-positive) and non-recombined (mTom-positive) acinar cells were distinguished by fluorescence.

### Intravital subcellular microscopy (ISMic)

Mice (18–34 g) were anesthetized by intraperitoneal injection of ketamine (100 mg/kg) and xylazine (15 mg/kg). Submandibular salivary glands or pancreas were surgically exteriorized and stabilized on a coverslip mounted on the microscope stage as previously described.^6^ Glands/pancreas were maintained with intact vasculature and innervation during imaging.

### β-adrenergic stimulation and inhibitor treatment

Secretory stimulation was induced by subcutaneous injection of isoproterenol (ISOP; 0.01 mg/kg). For inhibition experiments, exposed glands were bathed in 50 μM para-Nitroblebbistatin ^23^ (in 1.5% DMSO saline) for 10 min before stimulation. Control glands received vehicle alone. For perfusion experiments, solutions were applied at 10 μL/min using a programmable syringe pump.

### Intravital single-molecule microscopy (iSiMM)

Single-molecule imaging was performed on a Nikon Ti microscope equipped with a custom highly inclined and laminated optical sheet (HILO) illumination path using a 100× NA 1.49 oil immersion objective. Excitation was provided by a 488-nm laser expanded and focused onto the back focal plane of the objective. The incidence angle was optimized empirically to maximize signal-to-noise at imaging depths of ∼15–30 μm. Images were acquired at 30 ms per frame.

### iSIM imaging

iSIM imaging was conducted on Nikon Ti body base integrated with VT-iSIM (VisiTech International) module. Z-stacks of samples were acquired using a 100x oil immersion objective lens (Nikon NA 1.49) and sCMOS camera (ORCA-Quest; Hamamatsu). Lasers, cameras, stages, triggering, and microscopy are all controlled through Nikon NIS-Elements Imaging software. 488-nm laser was employed to excite GFP with a 525/50 bandpass filter.

### Spinning disk confocal microscopy

High-resolution 3D imaging was performed on a Nikon Ti2 spinning disk microscope. Z-stacks were acquired at 0.05–0.3 μm intervals with exposure times of 80–300 ms. Image alignment, denoising, and 3D deconvolution (Richardson–Lucy algorithm) were performed using Nikon Elements software.

### Image processing and analysis

Image processing was performed using Fiji/ImageJ ^30^, Nikon Elements, Imaris, and PRISM. Drift and motion were minimized by stabilizing the external gland as described before ^4,7,8^. When required, drift correction was performed using StackReg. Where required, Nikon Denoise.ai and Richardson–Lucy deconvolution were applied.

### Single-molecule detection and tracking

Emitter detection and trajectory reconstruction were performed using TrackMate^13^ with a Laplacian of Gaussian detector (estimated diameter 0.6 μm), sub-pixel localization, and median filtering. Tracks were generated using a simple LAP tracker (maximum linking distance 0.9 μm; maximum frame gap 2). Tracks shorter than four frames were excluded. Trajectories were exported to Spot-On ^2^ for two-state kinetic modeling (bound/free model, localization error fitted from data).

### ThunderSTORM reconstruction and localization-density measurements

Super-resolution density reconstructions were generated using ThunderSTORM^14^ from 10,000-frame datasets with default parameters and cross-correlation-based drift correction and reconstructions were displayed using average shifted histograms. Localization-density lengths were measured in Fiji along line profiles until signal intensity returned to baseline.

### Volumetric measurements

Acinar volumes were segmented manually using Imaris (v9.2.1) surface reconstruction tools based on membrane fluorescence. Surface volumes were exported and analyzed in PRISM.

### Basolateral membrane domain (BLD) width measurements

BLD width was measured using perpendicular line scans in Fiji. Width was defined as the distance between intensity plateaus at which mTom signal sharply decreased.

### NMII intensity measurements

Z-stacks (3 μm) were acquired to compensate for drift. Maximum projections were generated, and fluorescence intensity at BLDs was measured over time in Fiji. Data were normalized and analyzed in PRISM.

### FRAP analysis

FRAP experiments were performed using a 405-nm bleaching laser on the spinning disk system. Recovery was recorded for 8 min as 3-μm Z-stacks. Intensity was measured in Fiji and analyzed using easyFRAP ^31^ to determine half-times of recovery.

### Immunofluorescence

Cardiac perfusion was performed with PBS followed by 4% formaldehyde. Glands were cryoprotected in sucrose, embedded in OCT, and sectioned (10 μm). Sections were permeabilized (0.5% Triton X-100), blocked (10% NGS), and incubated with primary antibodies overnight at 4°C, followed by secondary antibodies. Slides were mounted in Fluormount-G.

### Tissue sample preparation and FIB-SEM

Mice were anesthetized and for stimulated group treated with ISOP. Salivary glands were surgically exposed, excised, washed in saline and fixed with Karnovsky’s fixative for 2 hours and then post-fixed in 2% osmium tetroxide with 1.5% potassium ferricyanide in 0.1 M sodium cacodylate buffer for 1 h at room temperature. Tissues were washed, stained overnight with 1% aqueous uranyl acetate at 4°C, washed again, and treated with lead aspartate at 65°C for 30 min. Tissues were dehydrated in graded ethanol (35%, 50%, 70%, 95%, and 100% x 5; 10 min each) and propylene oxide (PO; 100% x 5; 10 min each), infiltrated with Polybed 812 resin (resin: PO; 1:3, 1:2, 3:1, and 100%), embedded, and polymerized at 65°C for 48 h. Resin blocks were mounted on pin stubs, silver painted, and carbon coated (10 nm) before imaging in a Zeiss Crossbeam 550 FIB-SEM. Regions of interest were identified using a 1.5 kV, 1.1 nA SEM beam. FIB preparation followed the ATLAS3D workflow (Fibics, Ottawa, Canada), including a 1 mm platinum pad deposition, milling of tracking and autofocus lines before carbon coating, and trench milling with a 30 nA beam followed by 1.5 nA polishing. SEM images were acquired at 1.5 kV, 1.1 nA with 4 nm pixel size and 8 nm slice thickness using an in-lens EsB detector (grid 790 V). Image stacks were registered, inverted, and cropped in Atlas 5, then binned in 3dmod to generate 3D volumes at 8 nm isotropic resolution.

### Soleus transfection

Muscle electroporation was performed as described in DiFranco et al., 2009 following modifications^32^: For soleus electroporation, the mouse was anaesthetised with 3.5% isoflurane in O2 (Stinger Streamlight Rodent/Exotics table-top gas anaesthetic machine; Darvall Vet). The hindlimb was shaved with electric clippers and wiped with 70% ethanol to remove hairs and sterilize the skin surface. The mouse was placed on a heating pad to maintain body temperature and kept under anaesthesia during the transfection procedure via a rodent face mask. 40 µL of the plasmid (0.25 μg/μL solution in sterile saline; 10 μg/hindlimb) was injected onto the soleus muscle surface. 5 min after plasmid injection, electroporation paddle electrodes were placed on either side of the hindlimb, one electrode placed over the area where the DNA was injected and the second placed beneath the leg parallel to the first electrode with a distance of 4-4.5 mm. The voltage amplitude was adjusted to yield an electric field of ∼170 V/cm, then the hindlimb was electroporated with 6 pulses/20 ms interval/75 V (Square Wave Electroporation System - BTX, Harvard Apparatus; ECM 830). The mouse was returned to the cage for recovery. The optimal expression of the construct for intravital imaging was found to be 2-7 days post-transfection.

### Statistical analysis

A minimum of three independent animals and six cells per condition were analyzed unless otherwise indicated in the figure legends. For all analyses, biological replicates were defined at the animal level. Measurements obtained from individual cells were averaged per animal prior to statistical comparison. Comparisons were performed between animal means unless otherwise specified. Statistical analyses were conducted using Prism 10 (GraphPad Software). For comparisons between two conditions, unpaired two-tailed Student’s *t*-tests were performed on animal means. For comparisons involving more than two conditions, one-way ANOVA was performed on animal means, followed by appropriate post hoc testing where applicable.

Data are presented as mean ± SD unless otherwise stated. Statistical significance was defined as: ns, *P* > 0.05; *, *P* ≤ 0.05; **, *P* ≤ 0.01; ***, *P* ≤ 0.001; ****, *P* ≤ 0.0001.

## Acknowledgments

This work was supported in part by the Partnerships Program in the Office of Intramural Training & Education of the National Institutes of Health. R.W. and M.H. were supported by the National Institutes of Health, National Cancer Institute, Center for Cancer Research Intramural Research Program (ZIA BC 011682). K.N. and M.M. were funded in whole or in part by the National Cancer Institute, National Institutes of Health, under Contract No. 75N91019D00024. E.C.H. and P.W.G. were supported by grants from the Australian Research Council (DP160101623) and the Australian National Health and Medical Research Council (APP1100202, APP1079866).

We thank Dr. Renee Whan and members of the Katharina Gaus Light Microscopy Facility at UNSW Sydney for support with live rodent imaging, and Dr Elvis Pandzic for assistance with an earlier version of quantitative data analysis. The content of this publication does not necessarily reflect the views or policies of the Department of Health and Human Services, nor does mention of trade names, commercial products, or organizations imply endorsement by the U.S. Government. All schematics and cartoons were created using BioRender.

## Author Contributions

M.H. designed and performed experiments, analyzed data, and wrote the manuscript.

K.N. and M.M. performed and analyzed FIB-SEM experiments.

D.C. performed quantitative data analysis.

H.V and J. C. provided technical assistance with the execution of the experiments.

A.M. conceptualized and designed experiments and acquired funding.

T.M. conceptualized and designed experiments.

E.C.H., P.W.G. conceived and supervised the project, edited the manuscript, and acquired funding.

R.W. conceived and supervised the project and wrote the manuscript.

## Data availability statement

The data that support the findings of this study are available from the corresponding author upon request.

