## Supplemental Figures for "Intravital single-molecule microscopy reveals protein kinetics in live mammalian tissues"

**Content**

1) **Supplementary figures**
2) **Legend to supplementary movies**
1) **Supplementary figures**

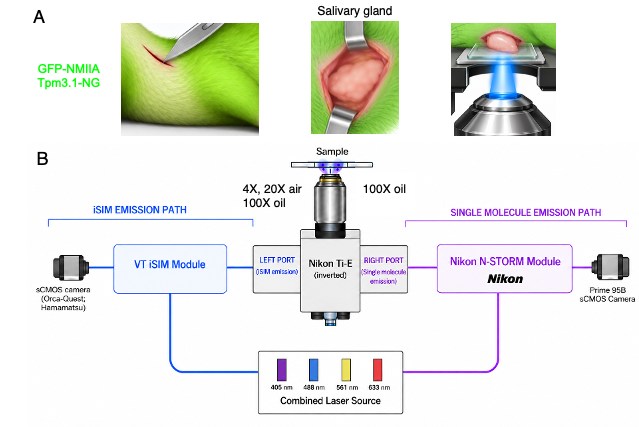

**Figure S1. Salivary gland preparation and optical configuration for intravital single-molecule microscopy.** (A) Schematic of the salivary gland preparation for intravital imaging in mice expressing GFP–NMIIA or Tpm3.1–mNeonGreen (Tpm3.1–NG). The submandibular gland is surgically exposed under anesthesia and positioned against a coverslip above the inverted microscope objective. The gland remains intact in the living animal, with its vascular and neural connections maintained. (B) Detailed optical layout of the Nikon Ti-E-based iSiMM microscope. The iSIM illumination and emission paths are shown in blue, and the single-molecule paths are shown in magenta. The combined laser source provides four available excitation wavelengths (405, 488, 561 and 633 nm), which are directed into two independent illumination paths using a polarization beam splitter. For iSIM, excitation is coupled through an optical fiber to the VT-iSIM module, where collimation and beam-expansion optics precede the excitation microlens array that generates multifocal illumination. Emission is collected through the same objective, directed through the left microscope port, passed through the collection optics and an emission microlens array that contracts the image twofold, and recorded on an ORCA-Quest sCMOS camera. For single-molecule imaging, excitation is coupled through a second optical fiber to the Nikon N-STORM/TIRF–HILO module, which controls beam size and incident angle for HILO illumination. Excitation is delivered through the quad-band excitation optics and a 4x air, 20x air or 100× oil-immersion objective. Single-molecule emission is collected through the same 100x oil-immersion objective, directed through the right microscope port, passed through the emission optics and quad-band filter, and recorded on a Prime 95B camera. The GFP–NMIIA experiments shown in Fig. 1 used 488-nm excitation.

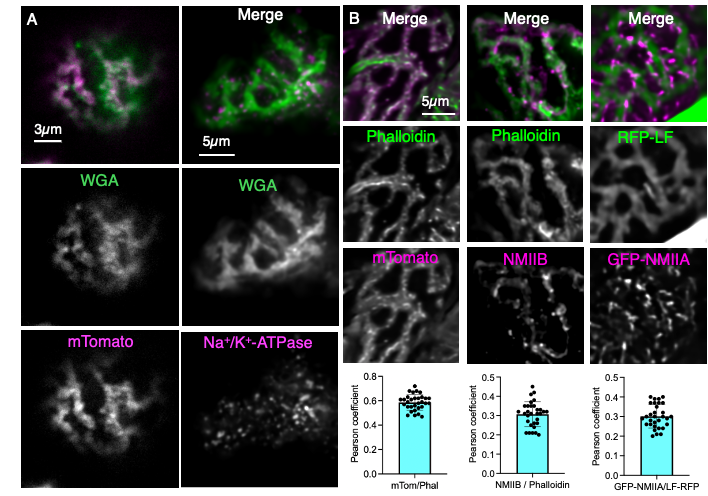

**Figure S2. Complementary membrane and cytoskeletal labeling supports the identity of basolateral membrane domains and their association with NMII. (A)** Fluorescence imaging of fixed salivary gland tissue using complementary membrane markers. Left, mTom-expressing glands labeled with WGA show overlap between membrane-targeted tdTomato (magenta) and WGA (green). Right, wild-type glands lacking the membrane reporter show overlap between WGA (green) and Na^+^/K^+^-ATPase (magenta), supporting the basolateral membrane identity of the labeled domains independently of mTom expression. Merged images are shown above the individual grayscale channels. Scale bars, 3 μm, left; 5 μm, right. **(B)** Association of F-actin and NMII with the basolateral membrane meshwork. Left, mTom-expressing glands labeled with phalloidin show overlap between the membrane reporter (magenta) and F-actin (green). Middle, NMIIB immunolabeling (magenta) and phalloidin staining (green) show the association of NMIIB with the actin-rich meshwork. Right, glands from GFP–NMIIA mice crossed with LifeAct–RFP mice show the spatial relationship between GFP–NMIIA (displayed in magenta) and LifeAct–RFP-labeled F-actin (RFP-LF, displayed in green). Merged images are shown above the individual grayscale channels. Graphs show Pearson’s correlation coefficients for each marker pair. For each comparison, *N* = 3 mice and 25 cells; data are mean ± SD. Scale bar, 5 μm, left.

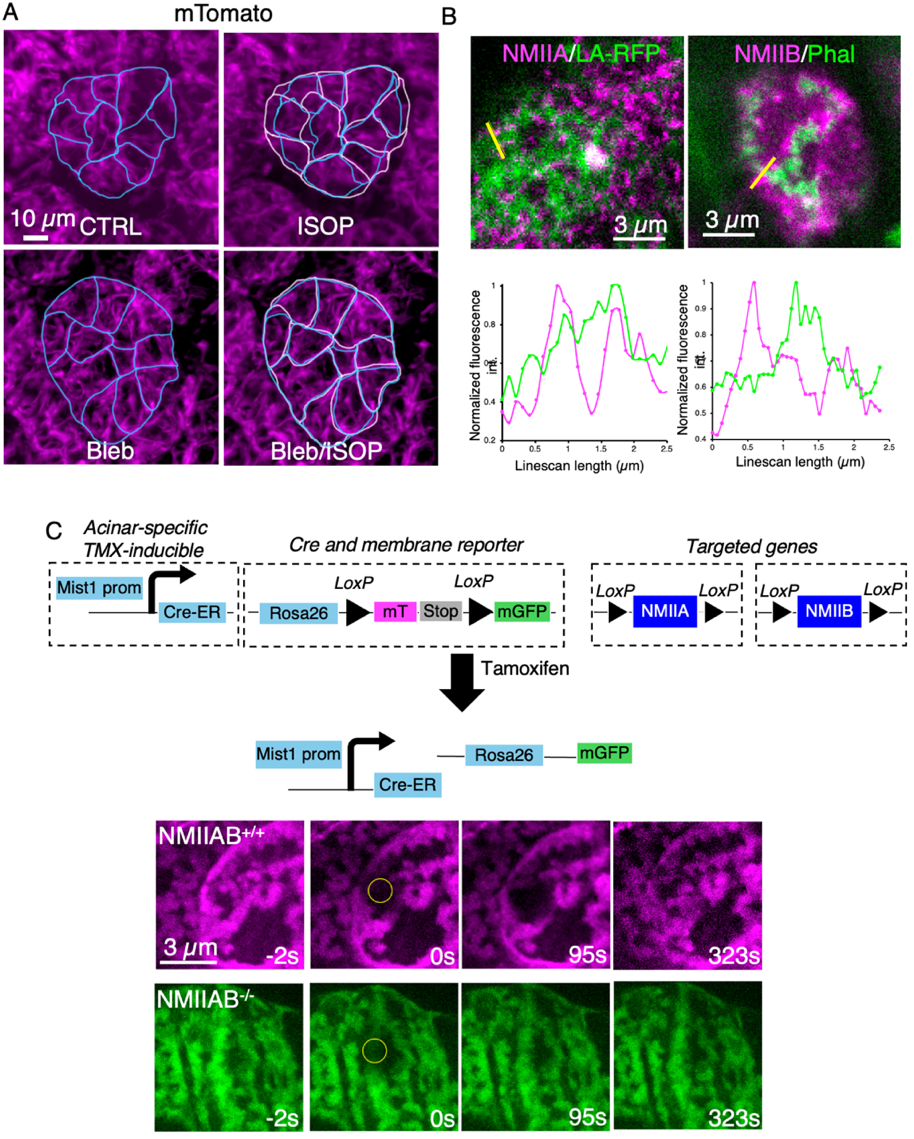

**Figure S3. NMII activity contributes to basolateral membrane organization and acinar cell expansion.** **(A)** Representative intravital images of salivary gland acini from mice expressing membrane-targeted tdTomato (mTom; magenta), under control conditions (CTRL), after isoproterenol stimulation (ISOP), after para-nitroblebbistatin treatment (Bleb), and after combined inhibitor treatment and stimulation (Bleb/ISOP). Light-blue and pink outlines indicate boundaries before and after stimulation, respectively. The images illustrate ISOP-induced acinar expansion and its attenuation after NMII inhibition, complementing the volumetric and BLD-width measurements in Fig. 3A. Scale bar, 10 μm. **(B)** Spatial relationship between NMII and F-actin within basolateral membrane domains. Left, GFP–NMIIA and LifeAct–RFP; right, NMIIB and phalloidin-labeled F-actin. NMII and F-actin are displayed in magenta and green, respectively. Yellow lines indicate the positions of the line scans, with normalized fluorescence intensity profiles shown below. Scale bars, 3 μm. **(C)** Conditional NMIIA/B knockout strategy and representative intravital fluorescence recovery after photobleaching (FRAP) experiments. Top, schematic showing the acinar-specific, tamoxifen-inducible Mist1CreERT2 driver, floxed NMIIA/B alleles and mTom/mGFP membrane reporter. Tamoxifen-induced recombination couples NMIIA/B deletion to a switch from membrane-targeted tdTomato (mTom) to membrane-targeted GFP (mGFP), allowing recombined NMIIA/B-deficient cells to be distinguished from neighboring non-recombined controls. Bottom, representative FRAP image sequences from mTom-positive control cells (magenta, upper row) and mGFP-positive NMIIA/B-deficient cells (green, lower row). Images show the membrane reporters before bleaching (−2 s), immediately after bleaching (0 s), and during recovery (95 and 323 s). Yellow circles indicate the bleached regions. Corresponding measurements of basolateral membrane domain width and fluorescence recovery half-times are shown in Fig. 3E. Scale bar, 3 μm.

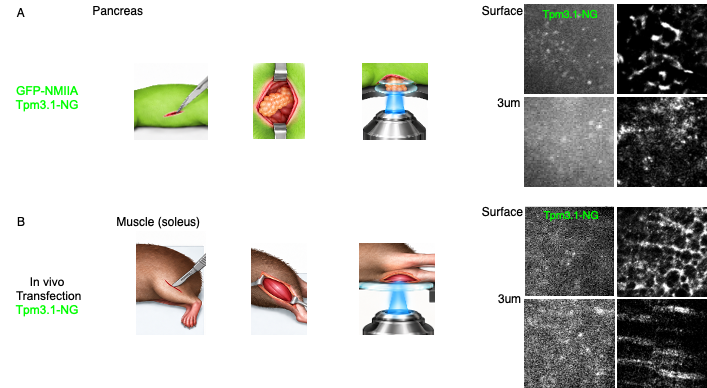

**Figure S4. Feasibility of single-molecule detection in pancreas and skeletal muscle using fluorescent reporter mice and in vivo transfection. (A)** Pancreatic preparation and imaging configuration for intravital single-molecule microscopy in fluorescent reporter mice. Schematics illustrate the location of the pancreas, its exposure for imaging, and positioning of the tissue against a coverslip above the inverted microscope objective while remaining connected to the living animal. The preparation is illustrated for mice expressing GFP–NMIIA or Tpm3.1–mNeonGreen (Tpm3.1–NG). Representative images on the right show Tpm3.1–NG fluorescence at the exposed tissue surface (top row) and 3 μm below the surface (bottom row). **(B)** Skeletal muscle preparation and imaging configuration following in vivo transfection with a construct encoding Tpm3.1–NG. Schematics illustrate the location of the soleus muscle, its exposure, and positioning against a coverslip above the inverted microscope objective for imaging in the living mouse. Representative images on the right show Tpm3.1–NG fluorescence at the exposed muscle surface (top row) and 3 μm below the surface (bottom row). Fluorescence images in both panels are displayed in grayscale.

2) **Legend to supplementary movies**

Movie S1.

Whole organ to single molecule in vivo imaging. Whole organ was imaged using iSIM and a 4x air, 20x air and 100x Oil lens, showing transition from lobules to acini. After a low background area was identified iSIMM shows NMIIA-GFP molecules on the basolateral membrane using the 100x Oil lens.

Movie S2.

Six-minute time-lapse imaging of a 2.5-µm Z-stack of the basolateral membrane of an acinus in a mouse expressing mTom (magenta). Z-stacks were acquired by spinning-disk confocal microscopy.

Movie S3.

Single optical planes extracted from a 2.5-µm Z-stack of the basolateral membrane of an acinus in a mouse expressing mTom (magenta). Basolateral membrane domains (BLDs) are visible as described in Fig. 1A. Z-stacks were acquired by spinning-disk confocal microscopy.

Movie S4.

Intravital single-molecule microscopy (iSiMM) time-lapse imaging of basolateral membrane domains (BLDs) in NMIIA–GFP and NMIIB–GFP mice. The left panel shows NMIIA–GFP molecules tracked by iSiMM; the right panel shows NMIIB–GFP molecules. Bottom panels display local molecular trajectories overlaid using TrackMate and color-coded by track duration, illustrating spatial confinement of molecular motion adjacent to BLDs. Experiments were performed in 3 NMIIA–GFP mice and 5 NMIIB–GFP mice.

Movie S5.

FIB-SEM volume of the basolateral membrane of acinar cells in CTRL and isoproterenol (ISOP)-stimulated mice. Following ISOP stimulation, the folded membrane architecture underlying BLDs is markedly reduced, as described in Fig. 3B and marked by arrows. Experiments were performed in 2 control mice and 2 ISOP-stimulated mice.

Movie S6.

Intravital single-molecule microscopy (iSiMM) time-lapse imaging of basolateral membrane domains (BLDs) in Tropomyosin3.1 (Tpm3.1-NG) mice. The left panel shows Tpm3.1 molecules tracked by iSiMM; the right panel displays local molecular trajectories overlaid using TrackMate and color-coded by track duration, illustrating Tpm3.1 binding to actin associated to BLDs. Experiments were performed in 3 Tpm3.1-NG mice.
